# Synaptic tagging and capture underlie neuronal co-allocation and temporal association memory in behaving mice

**DOI:** 10.1101/2023.09.17.558124

**Authors:** Y Sakai, B Brizard, J Zapata, X Zenelaj, A Tanti, V Camus, C Belzung, A Surget

## Abstract

Episodic memory has the ability to link distinct memories formed at temporal proximity (minutes-hours) into a coherent episodic representation. The neuronal mechanisms supporting such time associations remain however to be understood. The synaptic tagging and capture hypothesis (STC) provides a theoretical framework in which plasticity-related proteins produced for consolidating a memory trace at a synapse can potentially benefit to the consolidation of another trace at another synapse of the same neuron, thereby promoting neuronal co-allocations and temporal associations of memory traces. STC has however never been demonstrated in behaving animals, leaving its existence and functional relevance for memory formation unknown. We therefore investigated STC-like mechanisms in freely-behaving mice by recording hippocampal CA1 neurons during encoding and retrieval of distinct events. We found that reactivation of engram neurons at retrieval and the stability of place cells were strongly impaired by protein synthesis inhibition during encoding, but strikingly, were rescued in neurons that were coactive at another encoding close in time, having potentially benefitted from proteins produced at temporal proximity, as predicted by STC hypothesis. All our results together provide the first evidence of STC-like mechanisms in behaving animals and reveal an instrumental role of STC for time association of memory traces.

## Introduction

How are trivial events remembered, which would otherwise be forgotten, when they occur close in time to a memorable event? Episodic memory is indeed characterized by its ability to link independent events occurring at temporal proximity into coherent episodic representations ^1^.

Episodic memory traces (or engrams) are assumed to be supported by ensembles of active neurons (or “engram cells”) recruited in brain regions relevant for processing sensory information and integrating memory contents, such as the hippocampus ^2–5^. Distinct memories are typically encoded by well-separated, minimally overlapping ensembles of active hippocampal neurons ^5–8^. However, independent experiences may occasionally be co-allocated to shared groups of neurons when occurring close in time (from minutes to hours), resulting in memory traces with partially overlapping cell ensembles and, consequently, in memories that are likely to be recalled together ^9–14^.

The neurobiological underpinnings enabling such time-associated memories remain to be understood and characterized at a cellular scale. The synaptic tagging and capture model (STC) ^15^, by regulating long-term potentiation (LTP), provides a theoretical framework explaining how co-allocation of temporally proximal memories within shared neuronal ensembles might be achieved ^16,17^. LTP is a form of neuroplasticity crucial for memory formation, as it enables long-lasting enhancements of synaptic strengths between coactive neurons. According to the STC hypothesis, when a synapse of a neuron has received strong stimulations, it creates at the local site an early form of LTP (e-LTP) and a short-lasting “synaptic tag”, in addition to promote briefly the synthesis of plasticity-related proteins (PRPs). These PRPs can then be captured by the tagged synapse to durably consolidate LTP (l-LTP). Importantly, more moderate stimulations are them unable to trigger PRP synthesis on their own but can still create e-LTP and a tag. Without the presence of PRPs, both e-LTP and the tag decay after a few hours and LTP does not consolidate ^15^. However, a synapse tagged from moderate stimulations might potentially benefit from PRPs produced following strong stimulations at another synapse of the same neuron if they occur in relative temporal proximity (before the tag or PRPs decay). This heterosynaptic interaction might therefore lead transforming e-LTP into l-LTP even at moderately stimulated synapses, and theoretically enhance the proportion of co-allocated neurons for distinct memory traces encoded at temporal proximity ^15,18^.

Despite being behind many theoretical and neurobiological frameworks of memory, STC-like mechanisms have however never been demonstrated and validated in behaving animals, leaving their very existence and their functional relevance for learning and memory unknown and speculative. Indeed, STC-like heterosynaptic plasticity has only been demonstrated in brain slices *ex vivo* or suggested in anesthetized rodents ^19^ and by computational models ^20–23^. Indirect evidence for STC has also been proposed from behavioral studies in rodents and humans showing that memory performance is improved when learning is associated with a distinct novel experience close in time, the so-called “behavioral tagging” paradigm ^24–29^, but its link to STC-like mechanisms at a cellular level is uncertain.

In this study, we therefore aimed to evaluate in behaving animals the existence of STC-like mechanisms in conditions functionally relevant for memory formation and to determine how linking temporally proximal memories may involve STC at a cellular scale. For this purpose, we recorded pyramidal neurons of the CA1, a hippocampal subregion crucial for episodic encoding and temporal associations ^30,31^, using *in vivo* Ca^2+^ imaging in freely-behaving mice. Mice were allowed to explore distinct novel environments for memory encoding in sessions where protein synthesis had been inhibited (or not), enabling us to test the STC hypothesis by assessing whether recorded neurons can individually rescue their activity at retrieval, and supposedly their memory trace, when they had potentially benefitted from proteins produced at temporal proximity during encoding, as predicted by the STC model. Our study emphasizes the rescuing of memory traces in line with the STC hypothesis and provides the first evidence for STC-like mechanisms in behaving animals during memory formation, and particularly in consolidating and retrieving two temporally proximal memory traces at cellular scale.

## Results

### Experimental setup and hypotheses

To investigate the existence of STC-like mechanisms *in vivo*, we imaged GCaMP6f-expressing CA1 pyramidal neurons using a miniaturized epifluorescent microscope (nVoke, Inscopix) in mice behaving and encoding memories close in time. For this purpose, transgenic Thy1-GCaMP6f mice ^32^ aged over 11 weeks (7 females; 10 males) underwent lens implantation into the dorsal CA1 region (Figure 1A-B). Following recovery, the implanted mice were daily trained to freely explore an open-field arena where small pieces of food reward were randomly scattered to promote constant exploration for two sessions of 20 minutes per day. Once active exploration was observed in both sessions for 3 consecutive days (usually after 1 to 3 weeks of training), the animals underwent the behavioral procedure designed to test the presence of STC-like mechanisms (Figure 1C). During this procedure, CA1 pyramidal neurons’ single cell Ca^2+^ activity was recorded while the mice were exploring a novel square arena (40 cm x 40 cm) in two distinct novel environments (Room A and Room B) with a 50-minute interval for 2 consecutive days (sessions A1 and B1 on day 1, sessions A2 and B2 on day 2; Figure 1C). A protein synthesis inhibitor, anisomycin (ANI, 150 mg/kg, sc or ip; see Methods), was administered 30 minutes before admission to room B on day 1 in order to prevent PRP synthesis for this session. The capacity of anisomycin to temporarily inhibits *de novo* protein synthesis has indeed been well-characterized over decades by targeting the 60s ribosomal subunit and then preventing peptidyl transferase activity ^33–35^. Protein synthesis is inhibited in the mouse hippocampus in 30 minutes following systemic administration (150 mg/kg, sc or ip) and returns to baseline in less than 12 hours ^35^. Anisomycin has therefore been a useful tool for research, employed in various fields of biology to investigate the role of protein synthesis including in neurophysiological processes and memory formation ^36^.

**Figure 1.**
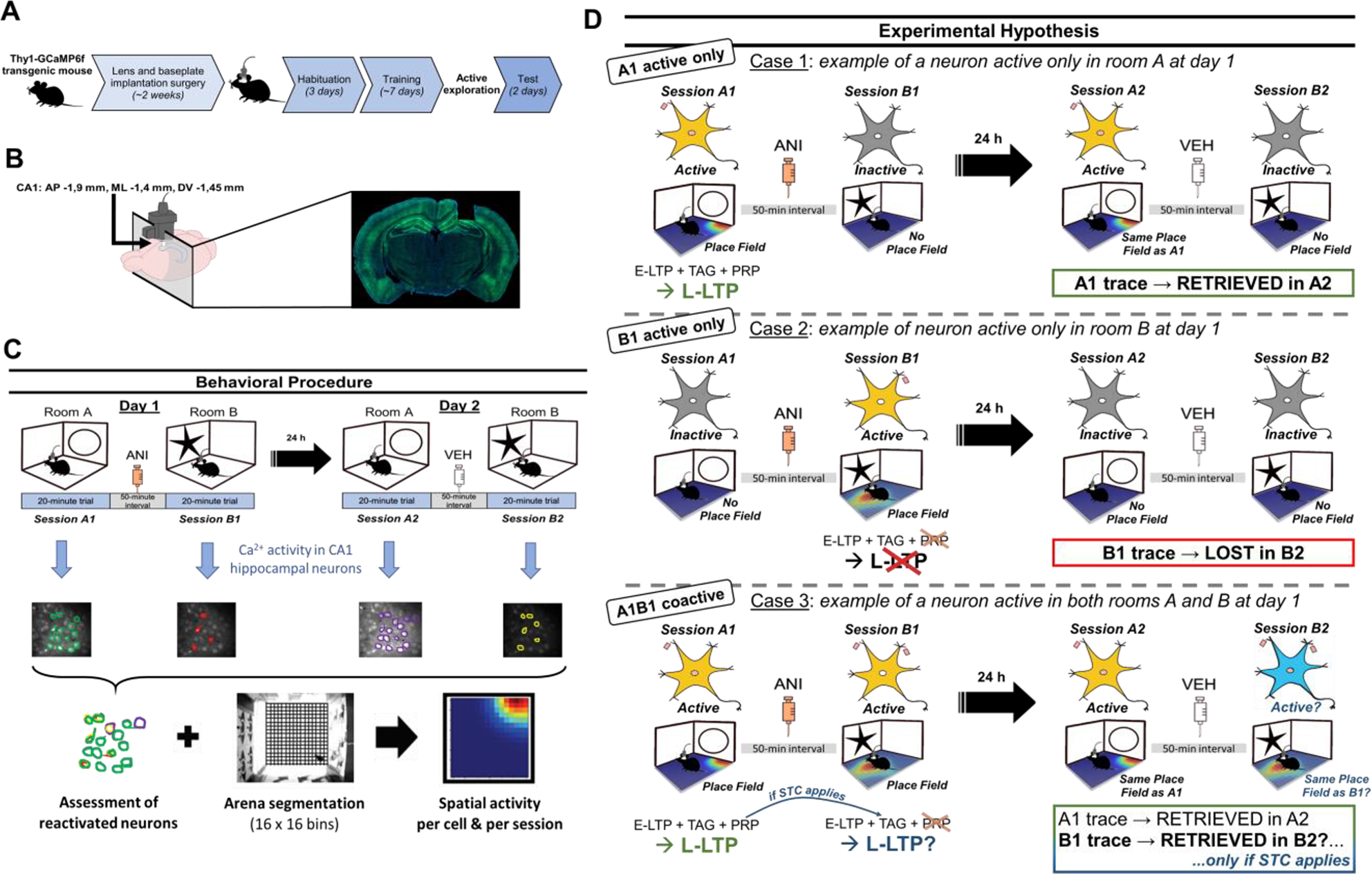
Schematic representation of the experimental outlines and hypothesis. (A) Timeline of the experiments. The baseplate and lens were implanted in the hippocampal region CA1 of Thy1-GCaMP6f mice aged over 11 weeks. After recovery and habituation, the animals were trained to explore the open field for two sessions (each one is 20 minutes). When active exploration was observed, neural activity was recorded while they were exploring a novel test arena. (B) Schematic image of lens implantation location. The lens was targeted to be implanted into dorsal CA1. The bottom panel represents a picture of an experimental mouse with a successful implantation into dorsal CA1, as evidenced by the trace of the lens in the brain. (C) The behavioral procedure. During the test, we observed CA1 neurons during exploration in four sessions spread across two days. Mice were exposed to two different rooms that have different environmental cues. On day 1, mice explored Room A then Room B (sessions A1 and B1), with the protein synthesis inhibitor, anisomycin (ANI, 150 mg/kg, sc or ip) being injected 20 minutes after A1 and 30 minutes before B1, in order to inhibit PRP synthesis for session B1. On day 2, mice explored the same Room A then Room B (sessions A2 and B2), with a vehicle (PBS 1x) injected between them. Ca^2+^ activity of CA1 neurons was recorded allowing identifying active “engram” neurons per session as well as probing their Ca^2+^ transients as a function of time. Ca^2+^ events of each neuron were aligned with the spatial exploration of the mouse within each session in order to determine their spatial activity and to identify place cells. (D) Schematic illustration of the experimental hypothesis that was used to test synaptic tagging and capture (STC)-like mechanisms at neuronal scale *in vivo* during memory encoding. Engram neurons active only in session A1 on day 1 are expected to be able to consolidate the trace encoded during A1 and to retrieve it during A2. Neurons recruited only in session B1 on day 1 are not able to produce PRPs because of ANI administration and, hence, are not expected to be able to consolidate the B1 trace and to retrieve it during B2. Neurons active in both sessions A1 and B1 on day 1 are expected to be able to consolidate the trace of A1 and also the trace of B1 (benefiting from PRPs synthesized during A1 according to STC hypothesis), and hence to retrieve both traces during sessions A2 and B2, only if STC applies *in vivo*.

For analysis, we compared across sessions neuronal activities and reactivation rates, as well as the stability of the spatial code of neurons identified as place cells ^37^ (i.e. neurons exhibiting selective activity based on the spatial location of the animal in its environment, delimitating one or several “place fields”, location(s) where the neuron is most active, assumed to provide information about animal’s location and experience). Our behavioral procedure was designed according to the following experimental hypothesis that would be valid only if STC-like mechanisms applied (Figure 1D). Among CA1 neurons that would have displayed activity on day 1, we might distinguish 3 main situations:

1. The case of a neuron that is active in session A1 but silent in session B1 (***A1 active only***). This neuron should be able to make e-LTP and a tag at the engaged synapse(s) in A1 and to produce PRPs that can convert e-LTP into l-LTP enabling trace consolidation and retrieval. These conditions should translate into the recruitment of the neuron in A2 (active in A2, silent in B2) and the conservation of its place field(s) in A2 if this neuron initially displayed place cell activity in A1 (*“A1 trace → retrieved in A2”*).
2. The case of a neuron that is silent in session A1 but active in session B1 (***B1 active only***). This neuron should be able to make e-LTP and a tag at the engaged synapse(s) in B1 but should not produce PRPs following anisomycin injection. Therefore, it should not be able to convert e-LTP into l-LTP, resulting in neither consolidation nor retrieval of the trace formed in B1. In consequence, the activity of the neuron in B2 should be fully independent from its activity in B1, resulting in either becoming silent in B2 or exhibiting a different spatial activity (*“B1 trace → lost in B2”*).
3. The case of a neuron that is active in both sessions A1 and B1 (***A1B1 coactive***). This neuron should be able to make e-LTP and a tag at the engaged synapses in both A1 and B1, and to produce PRPs during A1 only. The engaged synapse(s) in A1 should therefore be consolidated and an A1 trace should be retrieved in A2 (*“A1 trace → retrieved in A2”*). The neuron should not produce PRPs during B1, but the tagged synapse(s) during B1 should still benefit from PRPs already produced earlier during A1 according to the STC hypothesis. Accordingly, it should be possible to rescue the consolidation of the tagged synapse(s) in B1 and to retrieve the B1 trace despite the protein synthesis inhibition in session B1, but only if STC applies (*“B1 trace → retrieved in B2”*).

These conditions could potentially demonstrate the presence of STC-like mechanisms *in vivo* at cellular scale through a hypothetical differential response of “B1 active only” neurons (*trace lost in B2*) *versus* “A1B1 coactive” neurons during the session B2 (*trace rescued in B2*).

### Neuronal coactivation during encoding rescued anisomycin-impaired reactivation of engram neurons at retrieval

Here, we probed the presence of STC-like mechanisms through the proportion of CA1 neurons, initially active in either one or both sessions on day 1 (“A1 active only”, “B1 active only” or “A1B1 coactive”) that were also reactivated during retrieval on day 2 (Figure 2A). We defined a neuron as “active” in a session if the number of its detected Ca^2+^ events was above 5 in the session (see Methods) ^38,39^.

**Figure 2.**
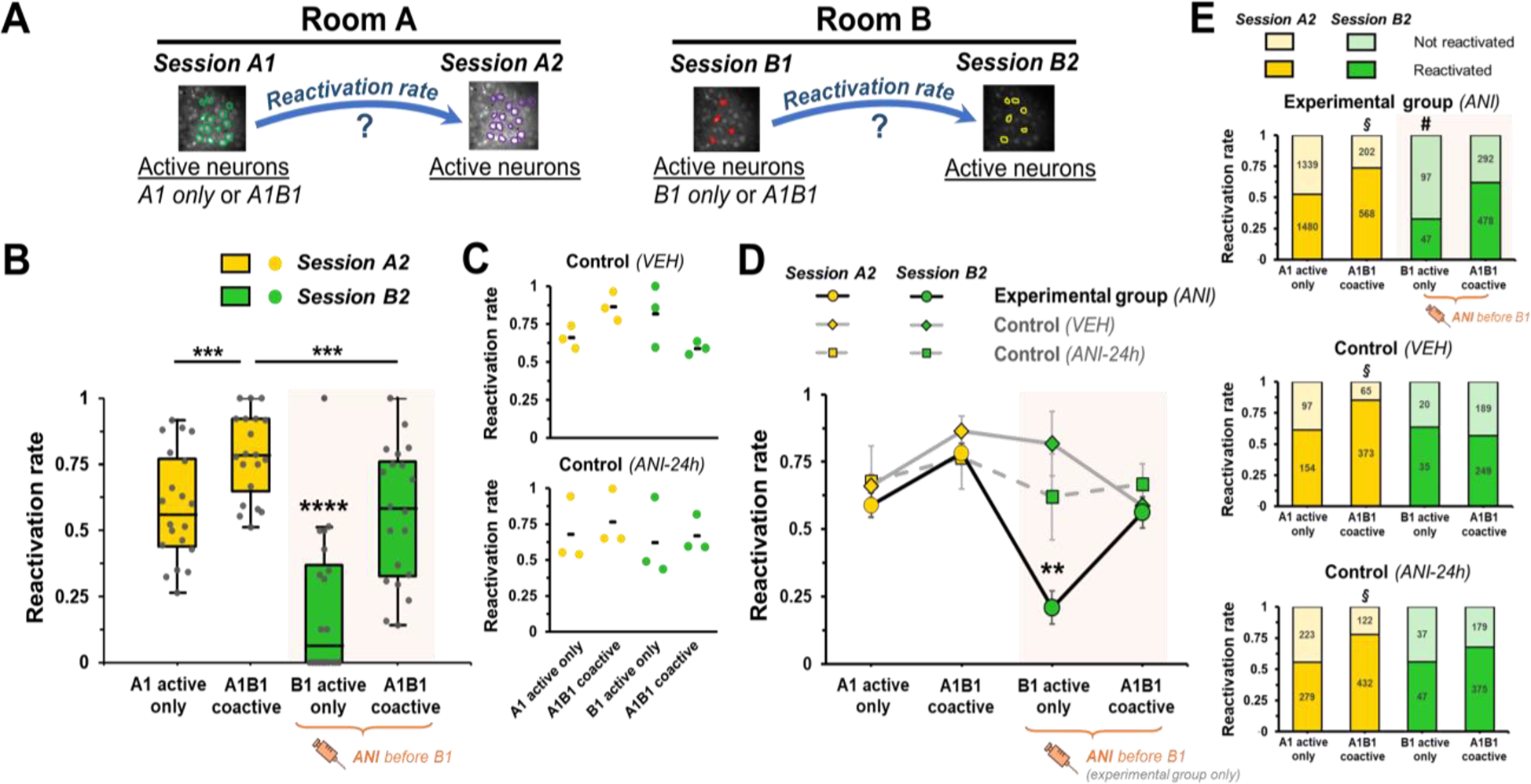
Neuronal A1B1 coactivation during encoding rescues anisomycin-impaired reactivation of B1 engram neurons at retrieval. (A) Schematic representation of the analytic approach based on evaluating the neuronal reactivation rates in each room on day 2 (session A2 *vs* B2) as an experimental readout. (B) Boxplot of the reactivation rate of engram neurons at retrieval (session A2 *vs* B2) according to their (co)activation status during encoding on day 1. ***, *p* < 0.01 between line-connected groups; ****, *p* < 0.0001 “B1 active only” neurons *vs* others. Gray dots indicate each individual mouse result (n = 20). (C) Neuronal reactivation rate of engram neurons at retrieval (session A2 *vs* B2) according to their (co)activation status for both control conditions: Control-VEH (upper panel, n = 3), Control-ANI-24h (lower panel, n = 3). Dots indicate each individual mouse result; Black dash indicates the mean. (D) Experimental condition effect on reactivation rate of engram neurons at retrieval in interaction with coactivation status and session. **, *p* < 0.01 “B1 active only” neurons (ANI) *vs* other neurons within condition and *vs* “B1 active only” neurons of other conditions (VEH and ANI-24h). Experimental group-ANI (n = 20), Control-VEH (n = 3), Control-ANI-24h (n = 3). (E) Frequency of neuronal reactivation based on the total numbers of detected cells depending on condition, coactivation status and session. #, *p* < 0.0001 “B1 active only” neurons *vs* other neurons within experimental group (ANI), and *p* < 0.05 “B1 active only” neurons *vs* same neurons in other conditions. §, *p* < 0.01 “A1B1 coactive” neurons in A2 *vs* other neurons in the same conditions. ANI syringe indicates neuronal groups that received an anisomycin injection (150 mg/kg, ip or sc) before encoding at session B1.

We found a significant effect of the retrieval session on day 2 (“A2” *vs* “B2”) in interaction with the coactivation status of the neurons on day 1 (“uniactive” *vs* “coactive”) (*F*_1,19_ = 5.834, *p* = 0.026; Figure 2B). Neurons that were active only in session B1 on day 1 (“B1 active only”), so following anisomycin injection, displayed a significantly lower rate of reactivation on day 2 (0.210 ± 0.061) compared to neurons that were active only in session A1 (“A1 active only”, 0.589 ± 0.046, *p* < 0.0001), highlighting that protein synthesis and hence PRPs are critical during encoding for consolidating and retrieving engram cell ensembles later. Indeed, with the inhibition of PRP production following anisomycin injection, we did not expect the B1 trace to be consolidated and retrieved by B1 active neurons later.

However, an exception might theoretically apply to the B1 active neurons that were also active in session A1, as these “A1B1 coactive” neurons might have benefited from PRPs produced during session A1, presumably still present during session B1 if STC-like mechanisms really occur during encoding. We indeed found a higher reactivation rate of “A1B1 coactive” neurons in session B2 compared to “B1 active only” neurons (0.562 ± 0.058 *vs* 0.210 ± 0.061; *p* < 0.0001), indicating that cell coactivation in both sessions on day 1 may rescue their incorporation into active cell ensembles for later retrieval in room B. This is presumably caused by PRPs produced during session A1, as neuronal activity during session A1 is the unique difference between “A1B1 coactive” neurons and “B1 active only” neurons, both recorded at the same time and compared within the same subjects.

In order to confirm the specificity of this outcome, our experimental group (ANI) was complemented with two control conditions where no differential response should apply according to our hypothesis: (1) a first experiment that involved a vehicle instead of anisomycin injection (Control-VEH, Figure 2C) to ensure that “B1 active only” neurons indeed exhibit similar reactivation rates as the other active neurons on day 1 when protein synthesis is not inhibited; (2) a second experiment that involved a 24h-interval between anisomycin injection and session B1 (Control-ANI-24h, Figure 2C) to ensure that the lowest reactivation rate of “B1 active only” neurons found in the experimental group (ANI) is specifically caused by the reversible, acute effect of anisomycin, and not by any other non-specific prolonged adverse effects that would impact delayed encoding and retrieval. In both control conditions, “B1 active only” neurons did not display lower reactivation rates (0.818 ± 0.118 in Control-VEH; 0.621 ± 0.159 in Control-ANI-24h) than any other classes of neurons (Figure 2C). When comparing experimental and control groups, we confirmed that only “B1 active only” neurons of the experimental group (ANI) displayed a significantly lower reactivation rate in session B2 (condition*session*coactivation interaction *F*_2,23_ = 5.834, *p* = 0.012; Figure 2D) compared to “B1 active only” neurons of the two control conditions (“B1 active only” ANI *vs* Control-VEH, *p* < 0.01, and *vs* Control-ANI-24h, *p* < 0.05). In addition, the reactivation rate in session B2 of “A1B1 coactive” neurons of the experimental group (ANI, 0.562 ± 0.058) was akin to those of Control-VEH (0.588 ± 0.025) and Control-ANI-24h (0.668 ± 0.075), indicating that neuronal A1B1 coactivation on day 1 was sufficient to rescue anisomycin-impaired reactivation rate in B2 at levels similar to those of control conditions. These results suggest that “A1B1 coactive” neurons may have benefited from protein synthesis and PRPs produced from neuronal activity during session A1 to consolidate the B1 trace and therefore plead in favor of the presence of STC-like mechanisms at cellular level during encoding.

Besides, it is worth mentioning that the reactivation rate of “A1B1 coactive” neurons in session A2 (0.782 ± 0.036; Figure 2B) of the experimental group, in addition to have been significantly higher than “B1 active only” neurons in session B2 as expected (0.210 ± 0.075, *p* < 0.0001), was also noticeably higher than “A1 active only” neurons in session A2 (0.589 ± 0.046, *p* < 0.01) and “A1B1 coactive” neurons in session B2 (0.562 ± 0.058, *p* < 0.01). However, this result may not be caused by the specific conditions of the experimental group (ANI) but rather be a regular feature as the reactivation rates of the same class of “A1B1 coactive” neurons obtained in session A2 of both control conditions exhibited comparable high levels of reactivation rates (Control-VEH, 0.864 ± 0.055; Control-ANI-24h, 0.765 ± 0.115; Figure 2D).

We also complemented our evaluation of the STC hypothesis using single cell activity as statistical unit and tested whether and how their reactivation might be explained by experimental conditions (“ANI” *vs* “VEH” *vs* “ANI-24h”), (co)activation status (“uniactive” *vs* “coactive”) and session at retrieval (“A2” *vs* “B2”). For this purpose, a log-linear regression analysis was performed allowing the evaluation of the relationships between these variables. The analysis reveals that the observed frequencies of neuronal reactivation were significantly dependent on the interaction between conditions, coactivation and sessions (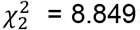; *p* = 0.012), an effect mostly caused by the lower frequency of “B1 active only” neurons (uniactive*session B2) observed in the experimental group (ANI) condition (frequency = 0.326) and confirmed by multiple pairwise post-hoc analysis (Figure 2E). Indeed, the observed reactivation frequency of “B1 active only” neurons of the experimental group (ANI) was significantly lower than all the other classes of neurons within the same condition (“A1 active only”, 0.525, *p* < 0.0001; “A1B1 coactive” in session A2, 0.738, *p* <0.0001, and in session B2, 0.621, *p* < 0.0001) as well as than the same class of “B1 active only” neurons of both control conditions (control-VEH, 0.636, *p* < 0.01; control-ANI-24h, 0.56, *p* < 0.05), confirming the results of our previous analysis. Again, the highest reactivation level of “A1B1 coactive” neurons in session A2 was confirmed and appeared to be independent of the experimental conditions (Figure 2E). Finally, the reactivation probability predicted by the log-linear regression model revealed that “B1 active only” neurons of the experimental group (ANI) is the unique neuronal class exhibiting higher probability of non-reactivation *vs* reactivation, with a predicted reactivation probability significantly lower than 0.5 (95% confidence interval [0.245, 0.434]), while A1B1 coactivation elevated the predicted reactivation in session B2 of the experimental group (ANI), despite anisomycin injection, to a probability significantly higher than 0.5 (95% confidence interval [0.568, 0.679]) and similar to all the other neuronal classes and all the other experimental conditions.

Altogether, these results demonstrate that A1B1 coactivation rescued the levels of anisomycine-impaired reactivation of B1 active neurons and support the existence of STC-like mechanisms *in vivo* during memory encoding.

### Neuronal coactivation during encoding rescued anisomycin-impaired spatial code of engram neurons at retrieval

We further investigated the presence of STC-like mechanisms through the examination of CA1 neurons identified as place cells. For this purpose, single cell Ca^2+^ activity was overlaid with the mouse trajectory in order to identify recorded neurons that displayed spatially tuned patterns of activity allowing them to qualify as place cells (Figure 3A). Active neurons were considered as place cells if the spatial information computed from their activity differed significantly from shuffled spatial information distributions (*p* < 0.05, see methods for details).

**Figure 3.**
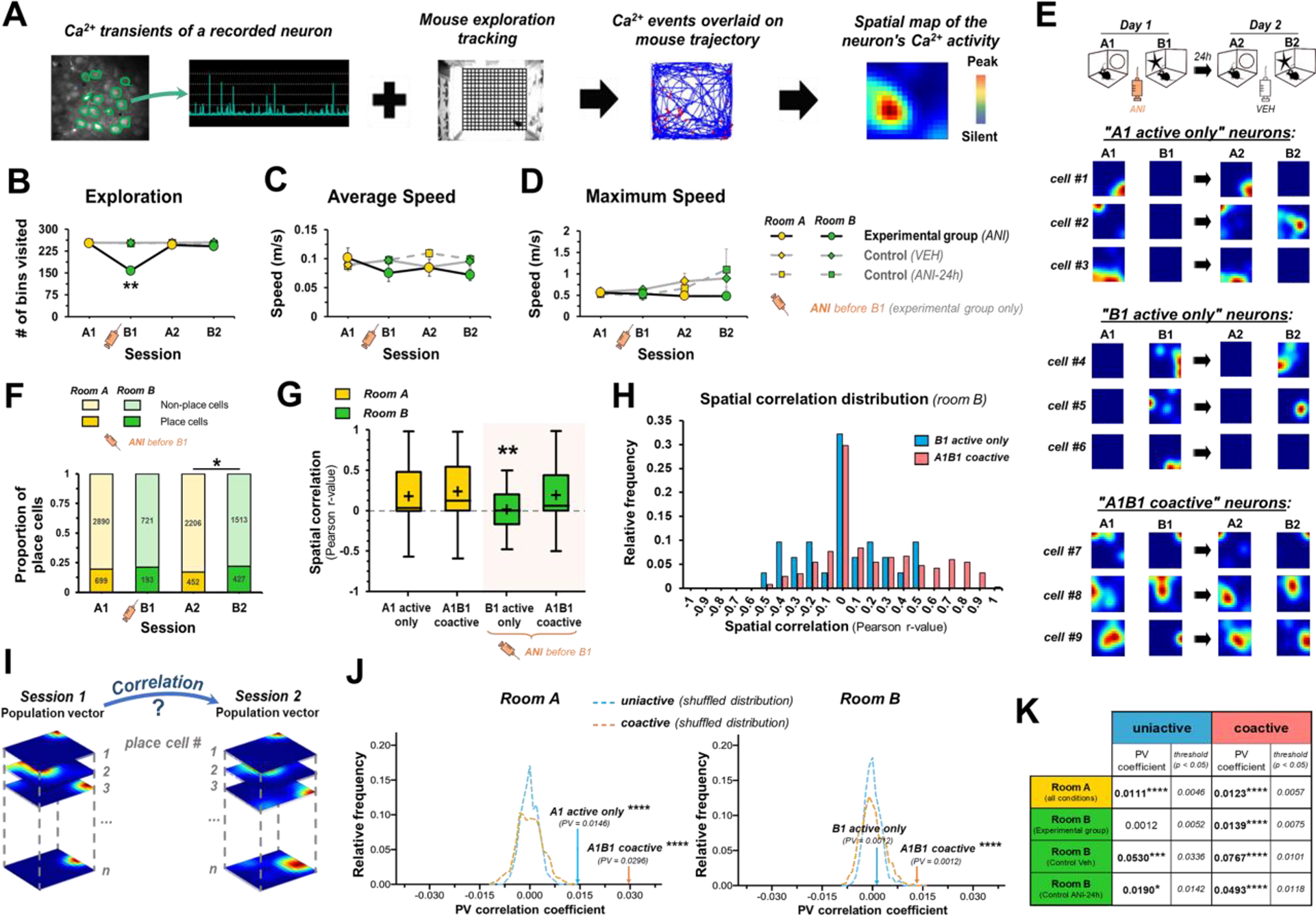
Neuronal A1B1 coactivation during encoding improves the stability of anisomycin-impaired spatially tuned activity of B1 engram neurons at retrieval. (A) Schematic representation of the workflow used to process behavioral and Ca^2+^ activity data. The color scale of the spatial map is from blue (silent) to red (peak number of Ca^2+^ events). Exploratory behavior and mobility of the mice were assessed through: (B) the number of bins the mice visited within the arena at each session, (C) the average and (D) maximum speeds the mice travelled during active bouts of each session. **, *p* < 0.01, B1-experimental group (ANI) *vs* other conditions in B1 and *vs* A1-A2-B2 sessions of the same condition. Experimental group-ANI (n = 20), Control-VEH (n = 3), Control-ANI-24h (n = 3). (E) Ca^2+^ activity spatial maps for each session (A1, B1, A2, B2) of representative recorded neurons identified as place cells and sorted according to their activation status on day 1 (“A1 active only”, “B1 active only” or “A1B1 coactive”). The color code was set as in (A) and was scaled dependently on each session and each cell. Spatially-tuned Ca^2+^ activity produced in session A1 was mostly retrieved in session A2 for “A1 active only” and “A1B1 coactive” place cells. Spatially-tuned Ca^2+^ activity produced in session B1 was mostly lost or uncorrelated in session B2 for “B1 active only” place cells while retrieved for “A1B1 coactive” place cells. (F) Numbers of identified place cells per session (n_A1_ = 699, n_B1_ = 193, n_A2_ = 452, n_B2_ = 427) of the experimental group (ANI) shown as relative frequency (f_A1_ = 0.196, f_B1_ = 0.211, f_A1_ = 0.170, f_B1_ = 0.220). 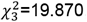, *p* < 0.001. B1 exhibited no significant difference with A1, A2 or B2. *, *p* < 0.05, A2 *vs* B2. (G) Boxplots of the spatial correlation coefficients of identified place cells of the experimental group (n = 1588) between sessions on day 1 and on day 2 according to the room (A or B) and their (co)activation status on day 1. ** indicates significant differences of “B1 active only” neurons *vs* all other groups: “A1 active only” neurons (*p* < 0.05), “A1B1 coactive” neurons in room A (*p* < 0.01) and in room B (*p* < 0.05). (H) Distribution of the spatial B1-B2 correlation coefficients of identified place cells in room B according to their coactivation status. Coactivation markedly increased the proportion of place cells with high spatial correlation coefficients (r > 0.5; 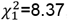, *p* = 0.004). (I) Schematic representation of the analytic approach for calculating population vector (PV) correlations between day 1 and day 2 within each room. The spatial maps of Ca^2+^ activity of identified place cells were stacked together into 256 PVs, one for each of the 16 x 16 bins segmenting the arena. The correlation between the PVs was computed between day 1 and day 2 and then compared to shuffled Ca^2+^ activity distributions. (J) The calculated PV correlations were significantly higher than shuffled Ca^2+^ activity distributions in room A for both “A1 active only” and “A1B1 coactive” neurons and in room B for “A1B1 coactive” neurons, but not for “B1 active only” neurons of the experimental group (ANI). ****, *p* < 0.0001 *vs* shuffled distribution. (K) Summary of PV correlations and statistical significances (*vs* shuffled distributions) according to experimental conditions. ANI syringe indicates neuronal groups that received anisomycin injection (150 mg/kg, ip or sc) before B1.

Anisomycin has been shown to transitorily reduce locomotion following injection through peripheral effects of protein synthesis inhibition ^40,41^. Considering that the second part of our analysis involved the spatial examination of cell ensemble dynamics and hence required a minimal exploration of the arena, we first assessed whether exploratory and locomotor activity of the experimental mice was affected by anisomycin. The arena was virtually segmented into 16 x 16 bins, resulting in 256 bins in total (Figure 3A). The number of bins the mice visited was significantly different across sessions and conditions (condition*session *F*_6,96_ = 6.086, *p* < 0.0001, Figure 3B). Although the session B1 of the experimental group (ANI), for which anisomycin had been injected 30 minutes before its onset, exhibited a significant reduction (*p* < 0.01) of the visited bins (B1, 158.26 ± 12.05) compared to other sessions (A1, 251.57 ± 1.57; A2, 246.61 ± 5.31; B2, 242.52 ± 5.80) and to control conditions (B1-VEH, 253.67 ± 0.67; B1-ANI-24h, 253.00 ± 1.53), the experimental mice still explored more than two thirds of the arena on average in B1 and exhibited normal average and maximal speeds when traveling (Figure 3C-D), indicating that anisomycin did not affect the mouse’s capacity to move normally during its active bouts and afforded a level of exploration large enough (1) to examine spatial Ca^2+^ activity of the recorded neurons and (2) to identify place cells (Figure 3E). Indeed, even if the lower exploration reduced the number of recorded cells that qualified as active neurons and place cells, we still identified an important number of B1 active neurons (914) and B1 place cells (193) and the proportion of identified place cells in session B1 was not different from other sessions (A1, 0.196; B1, 0.211; A2, 0.170; B2, 0.220; Figure 3F), indicating that locomotor activity as well as anisomycin did not affect the occurrence of place cells among the active cell ensembles. In addition, the experiments were designed to test our hypothesis using relevant comparisons carried out from the same pool of B1-B2 active neurons (“B1 active only” *vs* “A1B1 coactive neurons” in room B), exposed to the same conditions within the same animals, hence allowing us to control any possible nonspecific effects on either behavior or neuronal activity, if any.

We then investigated whether neurons that qualified as place cells and that were coactive on day 1, and supposedly benefitting PRPs produced during A1 if STC applies, were still able to consolidate a stable spatial code of room B and to retrieve it on day 2 despite anisomycin-induced protein synthesis inhibition in B1. For this purpose, we probed the stability and retrieval of place cells’ spatial codes on day 2 by calculating the spatial correlation across sessions in room A or in room B of the Ca^2+^ activity of neurons identified as place cells, splitting them between those coactive on day 1 and those active only in A1 or in B1. We found significant effects on spatial correlation caused by room (“room A” *vs* “room B”, *F*_1,1587_ = 9.081, *p* = 0.003) and by coactivation status of the neurons on day 1 (“uniactive” *vs* “coactive”, *F*_1,1587_ = 11.087, *p* < 0.001) as well as a profound trend for an interaction between both factors (room*coactivation, *F*_1,1587_ = 2.950, *p* = 0.058; Figure 3G). In particular, “B1 active only” neurons displayed significantly lower spatial B1-B2 correlations (room B-uniactive, 0.014 ± 0.047) compared to spatial A1-A2 correlations of “A1 active only” neurons (room A-uniactive, 0.180 ± 0.013, *p* < 0.05), suggesting that protein synthesis inhibition during encoding impairs place cell consolidation and retrieval. Importantly, “A1B1 coactive” neurons exhibited higher spatial correlations in room B than “B1 active only” neurons (0.191 ± 0.017 *vs* 0.014 ± 0.047; *p* < 0.05), which resulted in a higher proportion of neurons exhibiting elevated spatial B1-B2 correlations (r_B_ > 0.5, 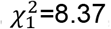, *p* = 0.004; Figure 3H). Accordingly, cell coactivation at both sessions of day 1 rescued the spatially tuned neuronal trace of room B at retrieval, suggesting that these cells benefited from PRPs produced during session A1, then endorsing the presence of STC-like mechanisms during encoding.

Populational vector (PV) analysis was also conducted for each room (Figure 3I). For this purpose, the spatial maps of Ca^2+^ activity of identified place cells were stacked together into 256 PVs, one for each of the 16 x 16 bins segmenting the arena. The correlation between PVs was computed between day 1 and day 2 for each room, each coactivation status and each condition. We then compared PV correlations calculated from our experimental data with shuffled Ca^2+^ activity distributions. In the experimental group, the calculated PV correlations were significantly higher than shuffled Ca^2+^ activity distributions in room A for both “A1 active only” (*p* < 0.0001) and “A1B1 coactive” neurons (*p* < 0.0001) and in room B for “A1B1 coactive” neurons (*p* < 0.0001), but not for “B1 active only” neurons (*p* = 0.618) (Figure 3J). The lack of PV correlation of the “B1 active only” neurons was not observed in both control conditions (VEH, *p* < 0.001; ANI-24h, *p* = 0.011; Figure 3K) confirming that the uncorrelated PVs of the “B1 active only” neurons of the experimental group ANI was indeed caused by anisomycin-induced PRP synthesis inhibition during B1.

In summary, we found for the experimental group ANI that (1) spatial Ca^2+^ activity associated with session A1 was mostly retrieved in session A2 for both “A1 active only” and “A1B1 coactive” place cells; (2) spatial Ca^2+^ activity associated with session B1 was mostly lost or uncorrelated in session B2 for “B1 active only” place cells, while it was retrieved for “A1B1 coactive” place cells (Figure 3E). Accordingly, neuronal A1B1 coactivation rescued anisomycin-impaired spatial codes of B1 active neurons at retrieval. Altogether, these results demonstrate that place cells could retrieve the spatial code of a first experience encoded under protein synthesis inhibition only if they had also exhibited activity at temporal proximity for another experience when protein synthesis was still possible. These conditions offered a temporal window for coactive neurons to benefit from PRPs and for consolidating both A1 and B1 traces, including the trace encoded under protein synthesis inhibition. Altogether, our results evidenced the existence of STC-like mechanisms at the neuronal scale in behaving animals during memory encoding.

### Spatial correlation in room A predicted the rescuing of anisomycin-impaired room B trace at retrieval

The STC hypothesis suggests that the production of PRPs in a neuron will be the determining factor for consolidation of its tagged synapse(s) and, by consequence, of the trace carried by this specified neuron. As a corollary, one may consider that the probability of reactivation and the level of spatial correlation between encoding and retrieval sessions would therefore be fostered by the level of PRPs produced during encoding. For instance, the level of spatial correlation in room A (A1-A2) in our experiments might increase in direct relationship with the level of PRPs produced at session A1. A prediction stemming from this STC corollary would be that, in our experimental group ANI, the probability for each “A1B1 coactive” neuron of rescuing its anisomycin-impaired room B trace would be associated with the level of PRPs produced during session A1, and therefore would be indirectly correlated with its level of spatial correlation in room A (for those identified as place cells). By contrast, such association and correlation might rather be marginal in control conditions (e.g. VEH and ANI-24h) where trace consolidation and retrieval can depend directly on the PRPs produced during session B1. To investigate this question, we therefore examined whether the level of spatial correlation in room A of “A1B1 coactive” neurons identified as place cells can indeed predict their probability of reactivation at session B2 and their level of spatial correlation in room B.

With this aim, logistic regression analyses were performed using spatial correlation as a predictor for reactivation of “A1B1 coactive” neurons identified as place cells. We first found, as anticipated, that reactivation probability in room A was increased as a function of the level of room A spatial correlation in all three conditions (experimental group ANI, 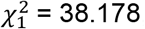, *p* < 0.0001; control VEH, 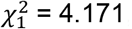, *p* = 0.041; control ANI-24h, 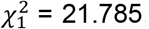, *p* < 0.0001, figures 4A-C). Of particular interest regarding STC hypothesis, the level of spatial correlation in room A was revealed to significantly predict the reactivation probability at B2 in the experimental group (ANI, 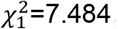, *p* = 0.006; Figure 4A). In a striking contrast to this group, reactivation probability at B2 did not relate to room A spatial correlation for both control conditions where PRPs can still be produced during session B1 (VEH, 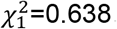, *p =* 0.424; ANI-24h, 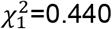, *p =* 0.507). On the other hand, spatial correlation in room B was found to be associated with reactivation probability at B2 but not at A2 for all three conditions as expected (experimental group ANI, 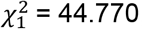, *p* < 0.0001; control VEH, 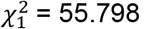, *p* < 0.0001; control ANI-24h, 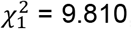, *p* = 0.002, supplementary figure S2). We therefore found a strict room-specificity of the association between spatial correlation and reactivation probability, with only one exception when PRPs cannot be produced (room B, experimental group ANI) but can still be provided by another experience of temporal proximity (room A, session 1). These results confirm the prediction made above and emphasize the existence of STC-like mechanisms *in vivo* during encoding.

**Figure 4.**
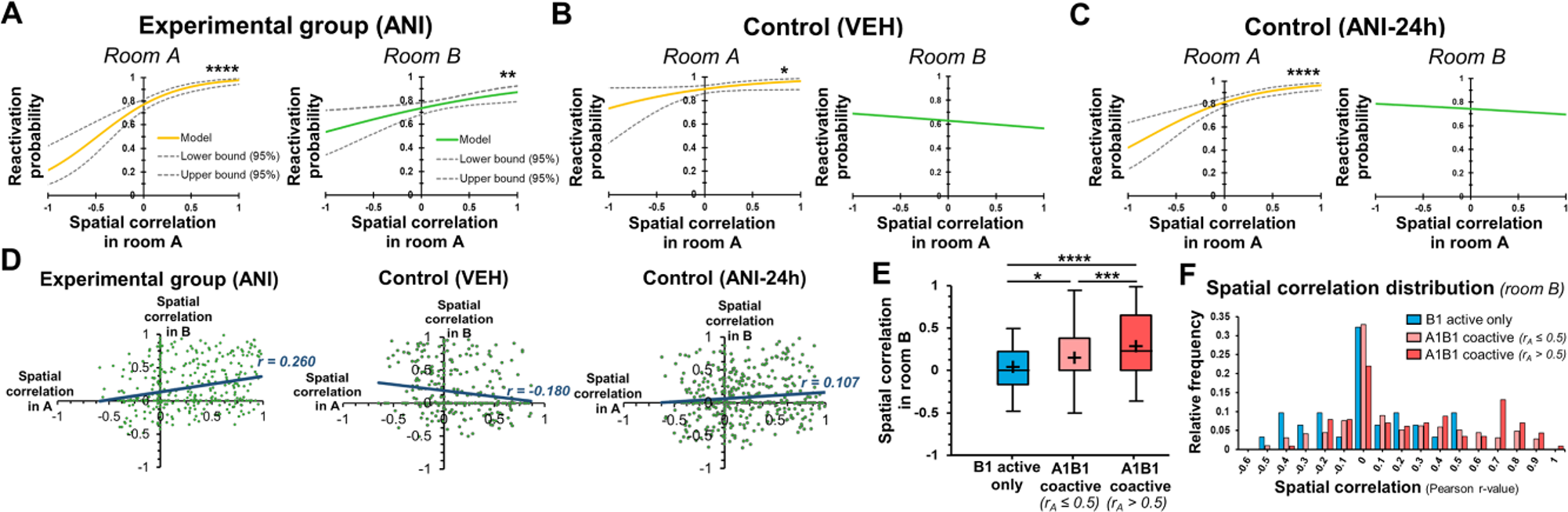
Spatial correlation in room A predicts the rescuing of anisomycine-impaired room B reactivation and spatial correlation of A1B1 coactive neurons in the experimental group (ANI). Reactivation probability of “A1B1 coactive” neurons identified as place cells was estimated by logistic regression using spatial correlation in room A as a predictor. (A) Reactivation probability of “A1B1 coactive” neurons of experimental group (ANI) were significantly enhanced by their level of room A spatial correlation in both sessions B2 (room B, green) and A2 (room A, yellow). Likelihood ratio test: *, *p* < 0.05; **, *p* < 0.01 and ****, *p* < 0.0001. (B – C) Reactivation probability of “A1B1 coactive” neurons of control conditions (VEH – ANI-24h, respectively) were significantly enhanced by their level of room A spatial correlation in session A2 (room A, yellow) but not B2 (room B, green). Likelihood ratio test: ****, *p* < 0.0001. (D) Room A spatial correlation significantly explained room B spatial correlation of “A1B1 coactive” neurons in interaction with experimental conditions (p < 0.0001). “A1B1 coactive” neurons of experimental group (ANI) displayed a significantly higher positive correlation (r = 0.260) between room A spatial correlation and room B spatial correlation than control conditions VEH (r = -0.180) and ANI-24h (r = 0.107), *p* < 0.0001 and *p* = 0.011, respectively. (E) Boxplots of the spatial correlation coefficients of identified place cells in room B of the experimental group (n = 434) between sessions on day 1 and day 2 according to their (co)activation status on day 1, “A1B1 coactive” neurons being split into two subgroups according to their room A spatial correlation level: r_A_ ≤ 0.5 and r_A_ > 0.5. A1B1 coactive neurons displayed significantly higher spatial correlation than “B1 active only” neurons, an effect significantly stronger for “A1B1 coactive” neurons displaying r_A_ > 0.5. *, p < 0.05; ***, p < 0.001; ****, p < 0.0001 between line-connected groups. (H) Distribution of the spatial correlation of identified place cells in room B according to their coactivation status and to their level of spatial correlation in room A. The proportion of place cells with high spatial correlation in room B (r_B_ > 0.5) is significantly increased by coactivation on day 1 and by high spatial correlation in room A (r_A_ > 0.5) (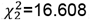, *p* < 0.001).

We then went further in addressing this question by assessing whether spatial correlation in room B of “A1B1 coactive” neurons identified as place cells can be explained by their spatial correlation in room A. We carried out an ANCOVA using (1) spatial correlation in room A and (2) conditions as explanatory variables for spatial correlation in room B. Our results revealed that room A spatial correlation significantly explained room B spatial correlation in interaction with experimental conditions (*F*_1,1168_ = 18.461, *p* < 0.0001). This interaction appeared to be mostly caused by the experimental group ANI that displayed a significantly stronger, positive association between spatial correlation in rooms A and B (ANI, r = 0.260, *p* < 0.0001) compared to control conditions (VEH, r = -0.180, *p* = 0.999; ANI-24h; r = 0.107, *p* = 0.026; ANI *vs* VEH, *p* < 0.0001; ANI *vs* ANI-24h, *p* = 0.011; Figure 4D). To illustrate and confirm this result, the “A1B1 coactive” neurons identified as place cells in the experimental group (ANI) were further divided into two subgroups based on their room A spatial correlation value (r_A_): (1) cells having a high room A spatial correlation r_A_ > 0.5 (“A1B1 coactive r_A_ > 0.5”) and (2) cells having a lower room A spatial correlation r_A_ ≤ 0.5 (“A1B1 coactive r_A_ ≤ 0.5”). They were then compared with “B1 active only” neurons for their spatial correlation in room B (*F*_1,431_ = 10.779, *p* < 0.0001; Figure 4E). Our results revealed that the cells that were coactive on day 1 and had high spatial correlation in room A (“A1B1 coactive r_A_ > 0.5”) presented a significantly higher level of spatial correlation in room B (0.285 ± 0.033) than “B1 active only” neurons (0.014 ± 0.047, *p* < 0.0001). The “A1B1 coactive r_A_ ≤ 0.5” cells also significantly differed from “B1 active only” neurons (p = 0.025) but at a much lower extent (0.153 ± 0.019) compared to “A1B1 coactive r_A_ > 0.5” neurons (0.285 ± 0.033, p < 0.001). This is confirmed by the distribution of room B spatial correlations, in which the proportion of place cells with high spatial correlation in room B (r_B_ > 0.5) was significantly improved by coactivation on day 1 and by high spatial correlation in room A (r_A_ > 0.5) (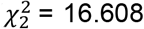, p < 0.001). Accordingly, “A1B1 coactive” neurons that highly retrieved their spatial code in room A also better retrieved their spatial activity in room B, but in the experimental group ANI only, when PRPs cannot be produced in room B on day 1.

Together, these results indicate that an “A1B1 coactive” neuron requires enough PRPs to be created in session A1 to rescue the consolidation and retrieval of its spatial code of room B despite protein synthesis inhibition at B1, and that this effect is much stronger and significant when the neuron was stimulated strongly enough to consolidate a strong memory of room A as well. This association between spatial correlation in rooms A and B was not found in control conditions, where PRPs can be produced directly from session B1. Indeed, control conditions represented conditions where consolidation and retrieval of room B trace did not depend significantly on the activity in session A1, and supposedly on the PRPs produced in A1 (in contrast to the experimental group ANI). Hence, these results corroborate those obtained using the “reactivation” analysis confirming (1) that protein synthesis and PRPs are critical for consolidating and retrieving spatial codes formed by hippocampal CA1 neurons during encoding and (2) that B1 traces can be rescued at retrieval, despite protein synthesis inhibition during B1, by coactivation in A1 at temporal proximity.

## Discussion

A fundamental feature of episodic memory is its capacity to link distinct memories formed at temporal proximity into a coherent episodic representation. Such temporal clustering is assumed to rely on heterosynaptic cooperation and cellular processes that would be capable of biasing neuronal allocation for different memory traces toward overlapping cell ensembles. STC has been proposed to provide such a mechanistic framework by potentially promoting neuronal co-allocation for time-associated memories within a defined temporal window. However, the presence of STC-like mechanisms had never been demonstrated in behaving animals and in conditions functionally relevant for memory formation to date. Our results demonstrate that trace retrieval at neuronal scale in the region CA1 of the hippocampus is impaired by protein synthesis inhibition during memory encoding but can be rescued for neurons that benefitted from proteins produced during another encoding at temporal proximity, as predicted by the STC hypothesis. Our study thus provides the first evidence of the existence of STC-like mechanisms at cellular level in behaving animals during memory formation.

The key rationale behind STC is that PRPs produced by an active neuron to consolidate a memory trace at synaptic level do not carry themselves any memory-related information and do not specifically target the tagged synapse(s) but spread all over the dendritic arborescence of the neuron and hence may also benefit other synapses that would have been tagged by other independent events at temporal proximity ^15,16^. Accordingly, blocking protein synthesis at specific timepoints during memory formation in different hypothesis-driven experimental setups allowed us to test STC-related predictions in behaving animals. Within this framework, our study found that: (1) active CA1 pyramidal neurons with no capacity to produce PRPs at memory encoding (e.g. “B1 active only” neurons) exhibited a severe decrease of reactivation probability and spatial correlation at retrieval, confirming that the availability of PRPs is essential during encoding to consolidate and retrieve memory trace at neuronal scale later. Prior activation in a first memory-creating context (e.g. room A) allowed CA1 pyramidal neurons that were also active in a temporally-close second context (e.g. room B, “A1B1 coactive” neurons), which occurred under PRP inhibition, to rescue their reactivation probability and their spatial code at retrieval for this second context, indicating that cells that are coactive for two temporally proximal events can utilize PRPs from the other event to consolidate both traces. (3) The level of spatial correlation in the first context where PRPs were available (e.g. room A) predicted the reactivation probability and the level of spatial correlation in the second context where PRPs were inhibited at encoding (e.g. room B), suggesting a possible quantitative relationship between PRP productions in room A and trace rescuing in room B in the experimental group (ANI). Taken together, these findings demonstrate that hippocampal CA1 neurons potentially benefit from proteins produced at a first event to rescue, consolidate and link another memory trace formed at temporal proximity, even under protein synthesis inhibition, as predicted by STC hypothesis. In consequence our study provides compelling evidence that STC-like mechanisms are at play and functionally relevant during memory formation in the hippocampus.

Other synaptic and cellular mechanisms have also been proposed to lie behind neuronal allocation to a memory trace ^16,42^. In particular, increasing neuronal excitability may also have an important role by enhancing the probability of a given neuron to be recruited and active during memory encoding ^43–46^. According to this view, neuronal allocation would not be random but predetermined by the level of excitability set up for each individual neuron before and during memory encoding. Hence, the subset of CA1 neurons that has been active for the encoding of a first event/context (e.g. room A) would exhibit an increase in excitability (lasting for hours or days), leading them to be preferentially recruited for a second event/context later (e.g. room B), fostering neuronal co-allocation to both events/contexts and promoting overlapping cell ensembles linking both memory traces ^10,47^. In addition, neurons with higher excitability levels at encoding would display significantly higher probability of carrying a memory trace and then of reactivation at retrieval ^43^. Might neuronal excitability hypothesis represent a potential alternative mechanism to STC, able to explain our results and particularly to underlie the rescuing of trace consolidation of room B in “A1B1 coactive” neurons? Considering this hypothesis in our experimental setups, the following prediction might be made: the probability of a neuron being recruited and active at session B1 as well as the levels of reactivation and spatial code stability at retrieval for both sessions A2 and B2 would increase as a function of the level of excitability settled at session A1. As a result, if neuronal excitability was the primary determinant of neuronal (co-)allocation in our study, it then would have implied that, in all three experimental conditions (ANI, VEH, ANI-24h): (1) “A1B1 coactive neurons” would represent the active neurons endowed with the highest levels of excitability in session A1 and therefore should exhibit the highest probability of reactivation at retrieval in both A2 and B2 compared to “A1 active only” and “B1 active only” neurons; and (2) a direct dependence of room B spatial correlation on room A spatial correlation should be observed for “A1B1 coactive neurons”. Accordingly, two distinct patterns of results can be anticipated depending on the mechanism that would be primarily at play for neuronal co-allocation in our study: a significantly higher reactivation at B2 for “A1B1 coactive” neurons compared to “B1 active only” neurons and a significant association between room A and room B spatial correlations of “A1B1 coactive” neurons, should be found: (1) for all three experimental conditions ANI, VEH and ANI-24h in the case of neuronal excitability hypothesis, or (2) only for the experimental group ANI in the case of STC. Our results indicate that the two expected results are well met in the experimental group ANI, but not in the control conditions VEH and ANI-24h, overturning neuronal excitability hypothesis and endorsing STC-like mechanisms as the major determinant of neuronal co-allocation and of trace B rescuing in our study.

Interestingly, the regulation of neuronal excitability has been shown to engage signaling pathways involving the calcium/calmodulin-dependent protein kinase II-IV (CaMKII-IV) and the transcription factor cyclic AMP-responsive element-binding protein (CREB), which have also been identified to trigger PRP synthesis ^48,49^. The exposure to a first memory-creating event therefore activates such signaling pathways leading to excitability increase in addition to PRP synthesis in the active neurons. In consequence, both hypotheses (i.e. STC and neuronal excitability) may well be two sides of the same coin, concurrently triggered. They may therefore represent complementary mechanisms for neuronal (co-)allocation and for linking time-associated memories but operating rather at different timescales from triggering events: a few hours for STC, one to two days for neuronal excitability ^10,47^. It would therefore be interesting to better understand the delimitation of the time windows that primarily involves STC *versus* neuronal excitability. It has been estimated from *ex vivo* electrophysiology that STC may be involved when intervals between sessions last no more than 4 hours ^18,50,51^. The unique study investigating STC *in vivo* in anesthetized animals only tested the 60-minute interval ^19^. In our study, we tested STC using 50-minute intervals between sessions (including 20-minute intervals before ANI injection and 30 minutes after the injection). It would be interesting to extend this interval in future studies in order to delimitate the maximum time between the two sessions where trace B can be rescued in our experimental conditions and therefore to determine the critical time window for a cell to utilize PRPs from previous activity in behaving animals and functionally relevant conditions.

In conclusion, using single cell in vivo Ca^2+^ imaging to monitor cell activity in behaving animals, our study provides the first demonstration of the existence of STC-like mechanisms at neuronal scale during memory formation, endorsing a critical role of STC in neuronal (co-)allocation to memory traces and consolidating theoretical frameworks on time-associated memories that built up on STC-like mechanisms.

## Method

### Animals

Seventeen C57BL/6J-Tg (Thy1-GCaMP6f) transgenic mice (Jackson Lab) aged at least 11 weeks at the beginning of the experiments were used in this project (male, n = 10; female, n = 7). The animals were group-caged until implantation was performed (see below). They were kept in cages (46 × 29 × 25 cm) enriched with tube and shelter under otherwise standard laboratory conditions (12/12 h light-dark cycle with lights on at 8:30 p.m. and room temperature at 22.2 °C). There was unlimited access to food and water until they were under food restriction protocols (see below). To reduce the use of animals, data were collected twice from the same mice as a general rule. Between the two tests, at least 2 days of training were carried out. All the procedure was approved by Directive 2010/63/EU guidelines on animal ethics (referral number: 2020112716449512-28454 approved by the ethical committee CEEAVDL).

### Surgery

Mice were first given an analgesic injection (buprenorphine 0.1mg/kg, sc, Buprecare®). After 10-20 minutes, the animals were placed into a gas induction chamber containing 2.5% isoflurane to induce anesthesia. When we observed no reflex from the animals, they were placed onto the stereotactic frame with their heads fixed. Anesthesia was maintained for the rest of the procedure using a mask, and the concentration rate was kept at 0.5-1.5% isoflurane. The flow rate and concentration were controlled in real-time to provide optimal anesthesia. After coordinating the Bregma and Lambda points, craniotomy was made either manually (1.6 mm diameter) or with a trephine drill (1.8 mm diameter) at stereotactic coordinates for lateral CA1 (AP-1.9mm, ML -1.4mm) at the center. Dura matter and the cortex above CA1 were removed by aspiration with a 30-gauge blunt needle while sterile phosphate-buffered saline (PBS, pH 7.4) was constantly applied to the exposed tissue. Following tissue aspiration, a baseplate with an integrated gradient refractive index (GRIN) lens (1 mm diameter, 4 mm length; ProView Integrated Lens, Inscopix) was implanted at AP-1.9mm, ML -1.4mm, DV - 1.45mm from the skull. Exposed tissue between the skull and the lens was covered with Kwik-Sil (WPI), and then Metabond dental cement was applied to cover the exposed area of the skull and to firmly attach to the baseplate. A 10% dermal Betadine at local, buprenorphine (0.1mg/kg, s.c), and Metacam (1mg/kg sc Provider) were applied as post-op procedure. Mice conditions were monitored twice per day for at least two days after the surgery, and the weight and their behavior were checked. For several mice (n = 12), GRIN lenses were not integrated with the baseplate. In those cases, the top of the lens was first covered with a Kwik-Cast (WPI) after lens implantation. The baseplate implantation was done at least 2 weeks after the first surgery. To set the optimal distance between the lens and the baseplate, the plastic base plate was attached to the miniscope (Inscopix), and then aligned parallel to the GRIN lens. The distance was adjusted manually to maximize the focus of GCaMP6f expressing cells for each animal. When the optimal field of view (FOV) was observed on the acquisition software (DAQ box, Inscopix), the miniscope was raised slightly higher to avoid over-shrinking of the dental cement. Metabond was applied to secure the baseplate and lens, and the plastic cap of the baseplate was then screwed on.

### Behavioral training and test

All behavioral experiments were conducted during the dark phase of the light cycle. The mice underwent habituation for 3 days at least 2 days after baseplate surgery or 2 weeks after integrated lens surgery. During this habituation, the mice were familiarized with handling by the experimenter as well as moving in the cage with the dummy scope mounted on their head. They were also habituated to food rewards (small pieces of biscuits or cereals). Food restriction was also started at the same time until the end of the test. All pellets were removed on the same morning as the start date of habituation, and only an adjusted amount of food was daily given to mice in the home cage (≤ 3.5g). The consumption of food each day and the weight of mice were monitored every day, and the weight of mice was kept at 85% of the weight from the initial body weight.

After habituation, the animals were trained to freely explore a square open arena (50 cm x 50 cm, masked) with the dummy scope (i.e. a factice scope with the same shape as the Inscopix nVoke miniscope) attached to their head during two sessions of 20 minutes. During the session, small pieces of food reward (i.e. biscuits, cereals) were scattered randomly and occasionally into the arena to encourage mice to explore. Between the two training sessions, 50 minutes intervals were used. Twenty minutes after the first session, saline was injected by sc (n = 11) or ip (n = 6) to replicate the test conditions, to habituate the mice to receive an injection and to reduce injection stress during the test. The mice were removed from the arena if they did not move from the same position for more than 4 minutes except for urinating and/or defecating, as we did not want to allow the mice to remain in the arena without actively exploring. For all sessions, we recorded the animal’s exploration to assess if the animal was exploring well enough. When we observed that they explored for 20 minutes twice without stopping for more than 3 days, the test was conducted. The arena was cleaned with 75% Ethanol and water after each session.

For the test sessions, the mice were attached to the miniscope and adjusted to an optimal FOV for each animal before starting the recording. The same recording parameters for each mouse were used for the rest of the test. The test was performed over 2 consecutive days, each containing two sessions with 50 minutes intervals between sessions (or 24 hours in a control experiment). In each session, the animals were allowed to freely explore a novel square open arena (40 cm x 40 cm, transparent) for 20 minutes. For session 1 and session 2, the arena was itself similar but different environmental cues such as visual cues, odors, noises, and recording rooms were used (room A for session 1, room B for session 2). To encourage the mice to explore, food rewards were given occasionally. Twenty minutes after the end of first session, the animals received anisomycin (150mg/kg, sc, n = 11, or ip, n = 6) or a vehicle (PBS 1x) in a control experiment.

All behavioral recordings were made by Media Recorder (Noldus) synced with nVoke recording software which was simultaneously recording the activity of CA1 cells activities. After each session, the arena was cleaned with 75% Ethanol and water. On the second day, the animal explored the same arenas using the same procedure, except that saline was administered instead of anisomycin.

### Drug treatment

Animals were injected with anisomycin (150mg/kg, Alomone), protein synthesis inhibitor, either by subcutaneously (n = 11) or intraperitoneally (n = 6). For ANI preparation, ANI was dissolved in PBS 1x using drops of 5% HCl to achieve the final concentration; pH was then adjusted to 7.4 using 4% NaOH.

## Data acquisition

Calcium imaging was performed using a one-photon miniaturized fluorescent microscope (nVoke; Inscopix, CA, USA) that can be mounted on the head of a mouse while the animal freely behaves. The data acquisition software (IDAS; Inscopix) was used at a frame rate of 20 Hz with LED intensities at 40 – 50 percent (0.8 - 1 mW) and gain at 1 – 1.5 that was optimized for each animal. Ca^2+^ imaging and behavioral recording were synchronized and triggered via TTLs to Media Recorder (Noldus) to align calcium transients and behavior. After 20 minutes from the start of the experiment, the data acquisition was automatically terminated.

## Data analysis

### Processing

Inscopix data processing software (IDPS; Inscopix) was used to process all the acquired Ca^2+^ imaging data (1440 × 1080, 20 fps). As the FOV was manually aligned for each day of recording by aligning with the blood vessel(s), all the video from the same test for each animal was registered as time-series which considered videos added to time-series as a list of the functions. Then the original Ca^2+^ imaging video was spatially downsampled by 4 and cropped at an optimal FOV for each animal. After pre-processing, a spatial filter and rigid motion correction were applied. Constrained non-negative matrix factorization for endoscopic data (CNMF-E) was used to identify regions of interest (ROIs). CNMF-E is one of the cell detection methods which is optimal for analyzing one photon microscopy data ^52^. All the detected ROIs were inspected manually to exclude cells that did not correspond to cells (i.e. ROIs with multiple components, fragments of a cell, and/or poor signal/noise quality). After Ca^2+^ signals were extracted from ROIs, the signals were deconvolved using a splitting conic solver (SCS). After denoising, the Ca^2+^ events were identified from Ca2+ transients (minimum size of 5 units of median absolute deviation (MAD) and at least a 200 ms minimal decay constant (τ)) and further inspected manually. All non-significant traces were zeroed.

### Place cell identification

We computed the spatial information calculation by Markus et al. (1994) using the unsmoothed-event-rate activity map of each cell.

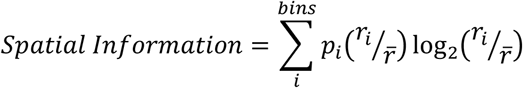

where *p*_*i*_ is the mouse’s probability of being in the *i* th bin; *r*_*i*_ is the rate of Ca^2+^ event in the *i*th bin; 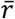 is the average of Ca^2+^ event rate in total; *i* is running over all the bins.

Then the statistical significance of spatial information was assessed by the shuffling method. The animal’s position during Ca^2+^ activity was shuffled 2,000 distinct times and we calculated the spatial information for each piece of shuffled data. Cells were considered as a significant place cell if it had more than 5 Ca^2+^ events per session, and the calculated non-shuffled spatial information was higher than 95% of shuffled spatial information (*p* < 0.05). For further analysis, we selected cells which were identified as a place cell in at least one of 4 sessions and had at least 5 events per session.

### Place field analysis

Behavioral data which was extracted from Ethovision XT (version 15 or 16, Noldus) and Ca^2+^ events were aligned and analyzed by custom Python code. The time when mice were not exploring was excluded (speed > 1cm/s). The square arena was divided into 256 bins (16x16, size 2.5cm x 2.5cm). For each bin, the occupancy time of the mice spent as well as the number of Ca^2+^ events were calculated. The rate map of a cell was calculated from Ca^2+^ events in a bin divided by total occupancy time in the bin. Then a Gaussian smoothing filter (σ=2 bins) was applied. The spatial correlation was calculated by Pearson correlation compared to the rate maps in two sessions in Room A or Room B. We then assessed the level of similarity in spatial representations from different sessions by Pearson correlation (Pearson’s *r*) using the smoothed rate activity maps. For each cell, we calculated Pearson correlations between the rate map vectors for corresponding sessions on each day to test the spatial map stability.

### Population vector (PV) correlation analysis

To assess the representations of each room across test days by the ensemble of the recorded hippocampal neuronal activity, we conducted PV correlation analysis ^37^. Cells which were defined as a place cell in at least one of 4 sessions and had more than 5 events per session were extracted from all the mice and concatenated as one. The unsmoothed rate activity of the whole cell population across animals in each bin was calculated, and the Pearson correlation between rooms on day 1 and 2 was computed. Next, we checked the statistical significance of PV correlation in each room by the shuffling method that the mice position during Ca^2+^ activity was shuffled 1,000 distinct times and calculated the PV correlation for each shuffled data. We then assessed if the PV correlation is higher than by chance using the value at 95% (after Bonferroni correction for multiple comparisons) of all shuffles data.

## Histology

At the end of the experiments, the animals received a lethal dose of pentobarbital (100 mg/kg, Dolethal®, Vetoquinol, Lure, France) via i.p. and were placed in their cage to avoid stress due to the change in the environment. When no reflex from the animal was observed, transcranial perfusion was performed. PBS with heparin was administered into the left ventricle through a needle, and the right atrium was incised to avoid overpressure. After blood was withdrawn, 4% paraformaldehyde (PFA) solution was perfused to fix the tissues. The head of each mouse was removed and placed into PFA for 2 nights to make sure of the fixation. After 2 nights, brains were harvested and placed in 10 % sucrose solution overnight, then placed in 20% sucrose solution and stored at 4 °C until sectioning. For histology, 40 µm coronal sections were made by cooled microtome (-20 °C, Leica CM 3050 S, Paris, France). All the brain slices were mounted onto the slides directly after cutting. Vectashield® mounting medium with DAPI (Vector Laboratories, Burlingame, CA, USA) was applied to the slides and then covered by a cover slip. Slides were stored at 4 °C. All the micrographs were taken by a Zeiss Z.2 Imager microscope in emitted-light mode using Zeiss ZEN software to check the expression of Ca^2+^ indicators as well as the location of lens implantation.

## Statistics

All the statistical analysis was performed using Statistica and XLStat. Explanatory variables that have been used in our analysis included for most of the tests: “session” (A2 *vs* B2) or “room” (A *vs* B), “coactivation on day 1” (uniactive *vs* coactive), condition (ANI, VEH, ANI-24h). When required following a significant effect including multiple experimental groups, we ran multiple planned comparison tests using Holm-Bonferroni corrections. All the details of the statistics, the tests used, and their results are mentioned in the legend and the manuscript and are fully provided in supplementary information.

## Notes

### Competing Interest Statement

The authors have declared no competing interest.

