## Supplementary Figures S1-S2 for "Synaptic tagging and capture underlie neuronal co-allocation and temporal association memory in behaving mice"

Sakai Y[1], Brizard B[1], Zapata J[2], Zenelaj X[1], Tanti A[1], Camus V[1], Belzung C[1], Surget A[1]
[1]UMR 1253, iBrain, Université de Tours, Inserm, CEDEX 1, 37032 Tours, France
[2]Inscopix Inc, 1212 Terra Bella Ave, Suite 200 Mountain View, CA 94303

### SUPPLEMENTARY FIGURES

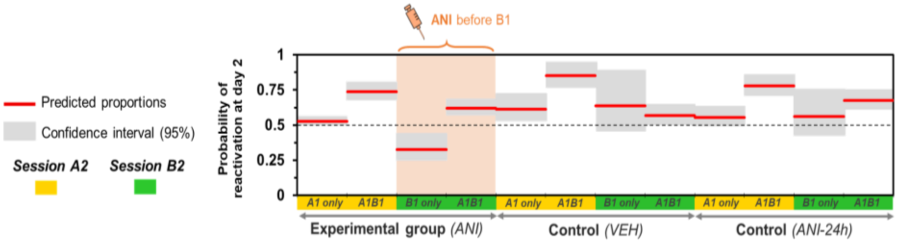


**Supplementary Figure S1. Probability of reactivation on day 2 according to condition, coactivation status on day 1 and session.**

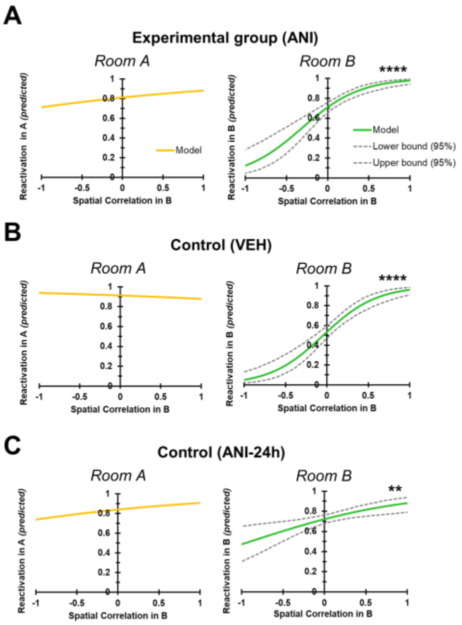


**Supplementary Figure S2. Reactivation probability of A1B1 coactive neurons identified as place cells was estimated by logistic regression using spatial correlation in room B as predictor.** Reactivation probability of A1B1 coactive neurons were significantly associated to their level of room B spatial correlation in B2 (room B, green) and not in A2 (room A, yellow), for (A) experimental group ANI, for control conditions (B) VEH and (C) ANI-24h. Likelihood ratio test: **, $p < 0.01$ and ****, $p < 0.0001$.
